# Epigenetic variation causes heritable variation in complex traits in the mollusk *Biomphalaria glabrata*, vector of the human parasite *Schistosoma mansoni*

**DOI:** 10.1101/2023.07.08.548202

**Authors:** Nelia Luviano, Marie Lopez, Fleur Gawehns, Bart Haegeman, Pierick Mouginot, Cristian Chaparro, Paola B. Arimondo, Benoit Pujol, Damien Pouzol, Céline Cosseau, Christoph Grunau

## Abstract

DNA methylation variation may play a role in phenotypic variation as it can be directly affected by the environment and be inherited. DNA methylation variations were introduced into the parasite vector snail *Biomphalaria glabrata* with low genetic diversity by chemical treatment in F0 and followed over 3 generations using epigenetic recombinant inbred lines (epiRILs). We observed phenotypic variation in complex traits such as fecundity and susceptibility to infestation by *Schistosoma mansoni* and DNA methylation differences in F3. Both, increase and decrease of infestation success (up to 100% and down to 20% prevalence in epiRILs and from 86% to 94% in control RILs) indicated variation in complex resistance/compatibility trait. Average prevalence in control RILs was 84±5% but only 68±21 % in epiRILs. Fecundity also changed and was in average 47±7% in control RILs and 59±18% in epiRILs, being 12% higher in epiRILs. We found that the heritability *h*^*2*^ of the fecundity in the epiRILs was between 0.5 and 0.6 depending on the method used to estimate it. We developed a model for introducing epimutant offspring snails into resident susceptible populations. If genetic assimilation of the resistant phenotype occured in a small fraction of the introduced epimutant snails, we predict that the susceptible phenotype is replaced by the resistant phenotype after 50-70 generations.

## Introduction

Epigenetic variation may play a role in phenotypic variation in animal populations as it can be affected directly by the environment and is likely to have higher spontaneous mutation rate than nucleotide sequences, and it may be inherited via non-mendelian processes (Chapelle and Silvestre, 2022). We define here epigenetic as heritable and reversible changes in gene function that are not based on changes in the DNA sequence and that can be mitotically and meiotically transmitted allowing for the inheritance of epigenetic marks across generations (Nicoglou and Merlin, 2017). We hypothesize that, genetic and epigenetic information are both part of an inheritance system, and both information allow for the generation of phenotypic variants on which selection can act. Inheritance conveys them between generations (Cosseau et al., 2016) with epigenetic heritability lower than genetic one. One of the most studied epigenetic marks is DNA methylation in cytosines leading to 5-methylcytosine (5mC). Environmental stress can induce epigenetic and plastic phenotypic changes and, after some generations, the phenotypic changes can appear without the environmental stress. However, it remains unknown by which mechanism the environmentally induced phenotypic change becomes genetically fixed (Nishikawa and Kinjo, 2018). The mutagenicity of DNA methylation patterns through the deamination of 5mC into thymine can potentially generate a final shift from epigenetic to genetic encoding (Danchin et al. 2019; Hanson and Liebl, 2022). Nevertheless, until now, there is no proof of such causal relationship between genetic assimilation and mutagenicity of DNA methylation marks. Therefore, in this work, we define the term “geneticassimilation” based on its heritability alone *i*.*e*. the part of variation that is similar between parents and offspring. We consider a trait as genetically assimilated if it acquired a heritability of 100% (h^2^=1) under constant environmental conditions.

In invertebrate species, there is evidence that DNA methylation is associated with developmental plasticity in the bee *Apis mellifera* (Elango et al., 2009) and it also plays a major role in the development and immune memory of the oyster *Crassostrea gigas* (Riviere et al., 2013, Fallet et al., 2022). DNA methylation is involved in behavioral plasticity in the bumble bee *Bombus terrestris* (Amarasinghe et al., 2014) and in the size variation of worker ants *Camponotus floridanus* (Alvarado et al., 2015). Its levels can influence alternative splicing in genes that are responsive to environmental stress, allowing to display transcript variability that in turn, will generate phenotypic variability (Flores et al., 2012). DNA methylation in intragenic regions is a common feature in some invertebrate’s species and in plants, and this type of methylation has been demonstrated to play a role in the generation of phenotypic variability mostly in plants (Seymour and Becker, 2017). In mollusks, despite being the second largest phylum of invertebrates, relatively little is known about the role of DNA methylation in generating heritable phenotypic plasticity. Given the ecological and sanitary importance of the phylum we wished to address this question with a focus on snail-borne diseases that are an emerging threat.

The mollusk *Biomphalaria glabrata* has a global DNA methylation pattern of the mosaic type (Adema et al., 2017) and 5mC occurs predominantly at CpG dinucleotides (Fneich et al., 2013). This snail is the intermediate host and vector of *Schistosoma mansoni*, a helminth parasite responsible for schistosomiasis, the second most important human parasitic disease after malaria, affecting about 230 million people worldwide. Not all *B. glabrata* snails are equally prone to infestation and prevalence in the field is actually very low. An important and complex trait is therefore resistance to infestation and/or host compatibility and knowledge of the molecular mechanism behind this phenotype would provide a rational for disease control. In *Biomphalaria* snails, DNA methylation shows responsiveness to parasite infestation by *S. mansoni* (Geyer et al., 2017; Knight et al., 2016). Furthermore, we previously demonstrated that DNA methyltransferase (DNMT) inhibitors induce phenotypic diversity in the morphometric traits of treated *B. glabrata* snails and its offspring (Luviano et al., 2021). We also showed that DNA methylation is essential for the snail (Luviano et al., 2023). Both, the existing knowledge on DNA methylation and phenotypic responsiveness to infestation make *B. glabrata* snails are an ideal system to study the role of DNA methylation in the generation of phenotypic plasticity. In addition, insights into the role of DNA methylation in this vector snail would be of practical importance for non-pharmacological control of schistosomiasis.

Here we tested whether the variation of DNA methylation plays a role in variability and/or plasticity of resistance. We used an integrative approach using chemical treatment, phenotyping, next generation sequencing and modelling to test our hypothesis. We used approaches that have been proven successful in model plants for the characterization of the contribution of DNA methylation to phenotypic variability. These approaches also permitted to disentangle the genetic and the epigenetic contributions to phenotypic variation. One of these methods relies on the generation of epigenetic recombinant inbred lines (epiRILs). In *Arabidopsis thaliana*, this approach allowed the discovery of epialleles contributing to phenotypic variation. This approach consists in reducing genetic diversity by crossing individuals that are genetically nearly identical but possess large DNA methylation differences. In *A. thaliana*, this was achieved by crossing wild-type plants with mutants of the DDM1 gene. The gene encodes an ATPase chromatin remodeler involved in the maintenance of DNA methylation and the mutant DDM1 plants exhibit a ∼70% reduction of global DNA methylation (Kakutani et al., 1995). Crosses with these mutant DDM1 plants introduced epigenetic diversity while keeping the genetic diversity low. The epigenetic recombinant inbred lines (epiRIL) were obtained by repeated backcrossing over six generations and subsequent self-fertilization for four generations (Johannes et al., 2009). The epiRILs realized in *A. thaliana* showed variation and high heritability in multiple phenotype traits in addition to stable inheritance of parental DNA methylation “epialleles” over eight consecutive generations (Roux et al. 2011). This epiRIL approach revealed the important role of demethylation in generating phenotypic variability for some complex traits and the stability of the epigenetic variation of these traits across multiple generations. Similar approaches have been performed in some animal species, e.g., the “epilines” obtained from genistein-injected eggs in the quail, (Leroux et al., 2017) but these approaches have, to our knowledge, not been applied to invertebrate species.

In order to test for the stable epigenetic basis of resistance to infestation and/or host compatibility in an invertebrate species, we decided to apply the epiRILs approach in the snail *B. glabrata*. This snail has a generation time of 70 days and can reproduce by cross-fertilization and self-fertilization. We treated the snails with DNMT modulators (DNMTm) to generate epimutations. Two new DNMTm were used: Flv1 (Luviano et al. 2021) and bisubstrate analogue compound 69 (Halby et al. 2017). For ease of readability, we will use “BA1” instead of “compound 69” in this work. Flv1 and BA1 treatments allowed us to generate snails with contrasting DNA methylation profiles: hypomethylated snail offspring of a cohort treated with Flv1, and hypermethylated snails that are offspring of those treated with BA1.We produced epimutant snails by producing crosses between hypo-and hypermethylated snails and pursuing the subsequent generation by self-fertilization. We then characterized the phenotypic variability of the snails regarding its fecundity and its response to the parasite infestation. As explained above we focused on prevalence and intensity of infestation that are among the most important phenotypic traits for parasite transmission, and fecundity is used as a proxy for fitness.

## Material and methods

### Ethics statement

*Biomphalaria glabrata* individuals of the Brazilian strain (*Bg BRE*) were used for this study. This strain of *B. glabrata* was maintained by inbreeding in our laboratory since 1975. First, parasite eggs were extracted from the liver of mice infested with *Schistosoma mansoni* Bresil (*SmBRE*). Then eggs were exposed to the light of a lamp, which induced hatching. Miracidia larvae were harvested with a pipette to choose 5 miracidia per snail for infestation experiments. The Direction Départementale de la Cohésion Sociale et de la Protection des Populations (DDSCPP) provided the permit N°C66-136-01 to manipulate on animals.

### DNMTm treatments

During previous work, treatments with a new flavanone-type DNMT inhibitor (Pechalrieu et al., 2020) named Flv1 were performed on the snail *B. glabrata* and showed high efficiency to reduce global DNA methylation in two consecutive generations (Luviano et al. 2021). In this work, we decided to use this flavanone compound and a second type of DNMTm compound 69 (Halby et al., 2017), called herein BA1. Both compounds have different mechanisms of action. Flv1 and BA1 were used to treat *B. glabrata* (Brazilian strain *BgBRE*) mature adults of 5-7 mm of shell diameter for 10 days. Stock solutions at 10 mM were made for each compound in ultrapure Milli-Q water and added to 1L of water (final concentration 10 μM) in an aquarium with 100 snails for each compound. Another aquarium without any compound was maintained as the control group with 100 snails.

### DNA extraction and global DNA methylation assays

The NucleoSpin® Tissue Kit (Macherey-Nagel, Düren, Germany) combined with the use of zirconia beads (BioSpec, USA, Cat. No. 11079110z) as described previously (de Lorgeril et al., 2018) was used for DNA extraction from whole body without shell of 30 *B. glabrata* snails per treatment group. The global 5mC of the 30 snails per treatment (F0 generation) and its offspring (F1 generation) was measured by the dot blot method based on the recognition of 5mC by anti-5mC antibody (Abcam, Cat. No. ab73938, Lot: GR278832-3) as described previously (Luviano et al., 2018).

The global 5mC level of the 4 control couples and the 15 couples with altered epigenomes used to raise the F2 generation were quantified using a commercial Kit ELISA Methylated DNA Quantification Kit (Colorimetric) (Abcam, ab117128) for detection of 5mC following manufacturer’s instructions. To quantify the absolute amount of methylated DNA, a standard curve was made by plotting the optical density values (OD) versus different concentrations of the positive control provided in the kit.

### Crossing experiment

A crossing experiment (Figure 1) was done with the offspring of the group treated with the compound Flv1, and the offspring of snails treated with the compound BA1. 15 couples were formed with one snail offspring of those treated with Flv1 and one snail offspring of those treated with BA1. We called these couples “epi-lines”. Another four couples with the offspring of control snails were formed and called “control lines”. For each control line and epi-line we put 3 snails into isolation that reproduced by self-fertilization and whose offspring was screened for divergent phenotypes. The 6 epi-lines with the most contrasting phenotypes were selected to realize a characterization of the phenotype with more statistical power, with 10 self-fertilization snails per epi-line and 10 snails in self-fertilization per control. After two generations of cross-fertilization between two individuals and one of self-fertilization (three generations) the F3 individuals were named epigenetic recombinant inbred lines (epiRILs) and control lines were named recombinant inbred lines (RILs), epiRILs and RILs correspond to the F3 generation (Figure 1).

**Figure 1.**
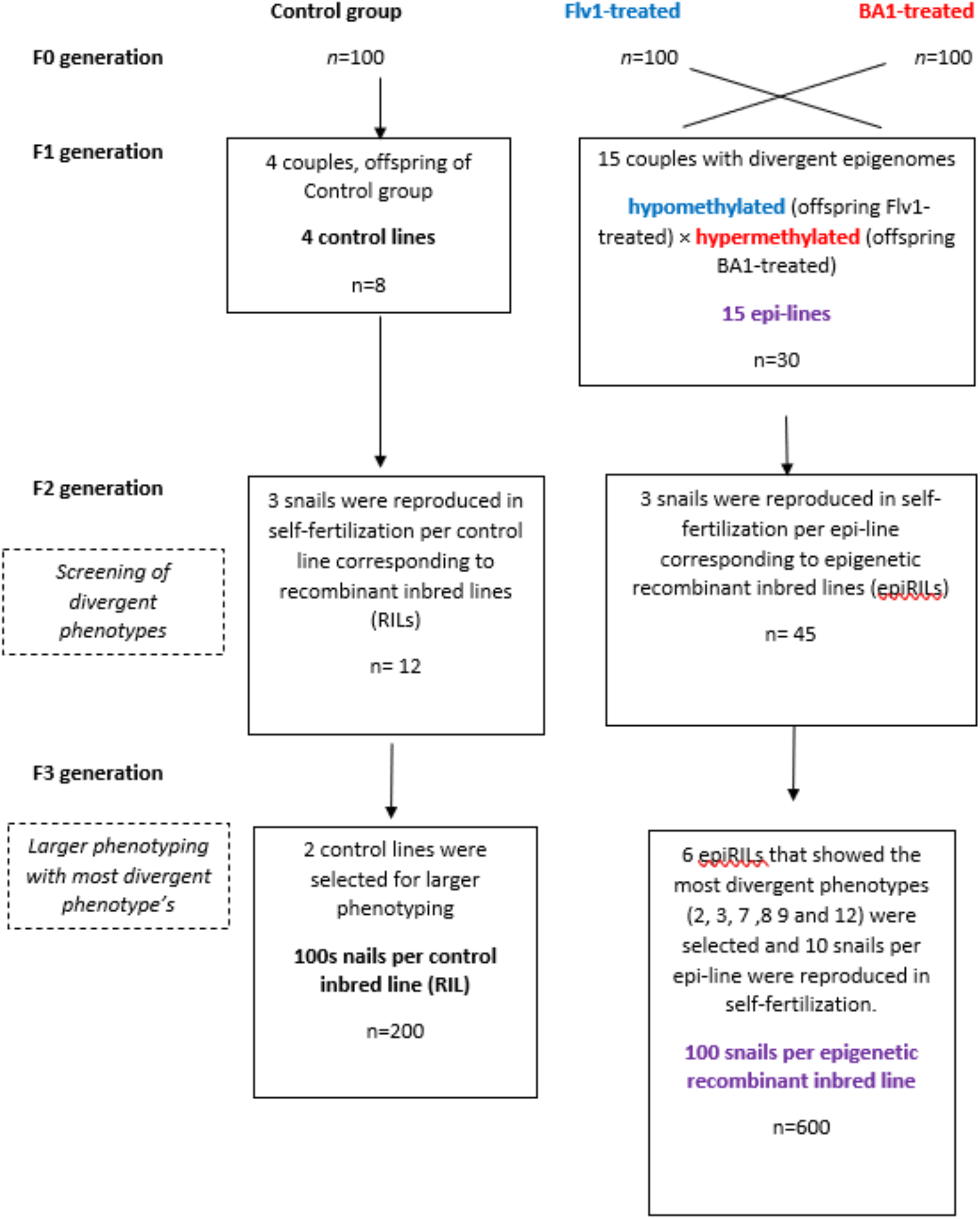
Experimental design of the multigenerational experiment. The snails of the F0 generation were exposed to the DNMTm Flv1 (in blue), BA1 (in red) or not exposed (control group, in black). In the F1 generation, four control couples and 15 couples with contrasting epigenomes (offspring of snails exposed with Flv1 and BA1, respectively) were formed. In the F2 generation, 3 snails were reproduced in self-fertilization for each control line and for each epi-line, 15 snails per epi-line and line of the F3 generation were used to measure their fecundity, the prevalence and intensity of infestation when exposed toS. mansoni. The 6 epi-lines that showed the most divergent phenotypes were selected for a second phenotyping with higher statistical power, 10 snails in self-fertilization by epi-line (named epiRILs) and 10 snails in self-fertilization for 2 control lines (named RILs).

### Snail phenotyping

Fecundity of the epi-lines and control lines was estimated by recording the number of eggs and hatched snails and calculating the hatching rate (number of hatched snails/ total number of eggs ×100). The most divergent phenotypes were screened on the basis of the 19 lines by keeping three snails per control line and epiline in self-fertilization (Figure 1). To measure prevalence and intensity of infestation, 15 snails per epiRIL and RIL were exposed to the parasite *S. mansoni*.

The two RILs and the six epiRILs with the highest fecundities in F3 generation were used to reach a large enough sample size for experimental infestations. Experimental infestations were done in the F3 generation RILs and epiRILs by exposing each snail to five parasite miracidia larvae. *S. mansoni* strain Bresil (*SmBRE*) was used in this study. The prevalence (number of infected snails / total number of snails exposed to parasite ×100) and the intensity of infestation (number of parasite larvae developed inside the snail) was measured per snail and then compared between control lines and epi-lines. In the second characterization of the phenotypes with more statistical power, with 10 self-fertilization snails per epi-line and 20 self-fertilization snails as controls, infestations were done in ten snails per RIL and per epiRIL (Figure 1).

### Reduced representation bisulfite sequencing by epiGBS

We applied an existing protocol of epinogenotyping by bisulfite sequencing (epiGBS) library preparation (Gawehns et al., 2022) to the snails characterized by significantly lower prevalence of infestation than controls. We modified some steps of the existing protocol. Instead of pooling all samples, we separated samples after the barcoding step. We also modified the bisulfite conversion step following the in-house protocol described in previous work (Luviano et al., 2021). Eight individuals of the epiRIL 8-7, eight of the epiRIL 9-3 and eight controls were sequenced. Paired-end sequencing (2×150 bp) was performed using Illumina Novaseq system at Fasteris (www.fasteris.com).

### Bioinformatics

We removed PCR duplicates, demultiplexed and trimmed the sequenced reads by using the most recent epiGBS2.0 snakemake pipeline (Gawehns et al. 2022). We took the Cutadapt (Martin, 2011) output trimmed reads and used the IHPE Galaxy interface (Afgan et al., 2018) to map the reads to the reference genome using BSMAP Mapper (Xi & Li, 2009). Single mapped reads were merged per sample and then merged by epiRIL (8 individuals per epiRIL). The merged mapped files were then used as input in the BSMAP Methylation Caller tool to get cytosine and thymine counts and then this file was used as input in the bsmap2methylkit tool to get tabular files that are used as input in MethylKit. We then used MethylKit (Akalin et al., 2012) to compare each epiRIL against the controls that were also merged. We used the parameters recommended to find differential methylated cytosines with weak methylation effects and small sample sizes that consist in a q value of 1 and 4% of methylation difference (Tran et al., 2018). We also performed SNP calling from bam files generated by BSMAP, for that purpose we used CgmapTools (Guo et al., 2018) that allows the SNP calling from from bisulfite sequencing data.

### Treatment effect on fecundity and heritability

The fecundity of the F2 generation descending from the F0 exposed to the DNMTm treatment was either inferior to the minimum fecundity of the control group (<35.7 eggs/offspring) or superior to the maximal fecundity of the control group (>40 eggs/offspring). Therefore, individuals of the treatment group were split in two groups: the higher and the lower fecundity groups. We did the same for the fecundity of the offspring generation. We test whether both the higher and lower fecundity groups are different from the control group. To do so, we built a linear model with parent and offspring fecundity as dependent variable and the fecundity expression group (control, higher, and lower) as explanatory variable. We checked the models for assumptions of linearity.

We estimated the heritability of fecundity by using a parent-offspring regression model. We calculated heritability with repetitive data in F2 generation (*n* = 45) by regressing with the offspring fecundity of epiRILs. We also regressed the median fecundity value of full sibs to avoid pseudo-replication in the regression analysis (*n*=17). The estimated heritability corresponds to the slope of the parent-offspring regression (Falconer & Mackay 1996).

### Mathematical simulation model

We developed a mathematical model to explore whether introducing epimutant snails can make a resident snail population more resistant. We assumed a constant prevalence for the resident snails, and a changing prevalence for the epimutant snails: low immediately after introduction and gradually increasing towards the prevalence of resident. The epigenetic variants are mostly not reversed in a predictable manner and are not as stable as genetic mutations (Pal and Miklos, 1999). As a result, epimutants and their offspring have a fitness benefit during the first generations after introduction, which can become persistent if the heritability of the epigenetic induced phenotype change in infestation prevalence equals the genetic heritability, which we called genetic assimilation in our model. To assess the effect of the introductions, we compared the average prevalence in the community of both epimutant and resident snails to the case without introductions. We considered different scenarios: a single introduction or repeated introductions, either without or with genetic assimilation.

A detailed model description together with simulation code is provided in the supplementary material. Here we give a brief overview: at each generation the model dynamics consisted of five steps. First, epimutant snails were introduced if the introduction scenario stipulated so. The number of introduced snails was set arbitrarily to 1% of the resident population. Second, snails were exposed to parasitic larvae, and each snail got infected with a probability equal to its prevalence. Prevalence differed between resident and epimutant snails, as described below. Third, snails underwent reproduction through self-fertilization and mortality, with a higher reproduction rate and a lower mortality rate for uninfected compared to infected snails. Fourth, the snail population was reduced to its original size, in order to keep the population size constant. Fifth, in the scenario with genetic assimilation, a small fraction of the epimutant snails (3% or 0.3% of the individuals per generation) underwent genetic assimilation. This was implemented by keeping their current prevalence value fixed for all following generations (i.e., heritability 1).

For the prevalence values, we associated the resident snails to the control group of the experiment and kept their prevalence constant, equal to *p*_*res*_ = 0.8. For the epimutants we set the prevalence immediately after introduction *p*^*0*^_*mut*_ = 0.3, a value observed for several epiRILs (see Results section). We used heritability *h* = 0.6, meaning that 60% of the epiRILs progeny are similar to their parents and to determine the rate at which this prevalence returns to the resident prevalence p_res_ (see Results section for the choice of the value). That is, the prevalence *p*_*mut*_*(n)* of epimutants n generations after 0n their introduction is given by *p*_*mut*_*(n) = p*_*res*_ *+ (p*_*mut*_ *– p*_*res*_*) h*^*n*^. The model analysis (see supplementary material) shows that the predictions are not dependent on the separate values of reproduction and mortality rate of infected and uninfected snails, but only on the fitness ratio of infected vs uninfected snails. We set this ratio equal to *W*_*inf*_*/W*_*uninf*_ = 0.7 based on earlier published results (Looker and Edges, 1979).

## Results

### DNMTm produce viable and fertile hyper- and hypomethylated snails in F1

The dot blot results showed that the Flv1 compound reduces the global 5mC% 2-fold in the F0 and F1 generation (Figure 2a-b, blue bars). BA1 reduces the global 5mC% 2-fold in the F0 generation but led to a 2-fold higher global 5mC% in the F1 generation compared to control (Figure 2a-b, red bars).

**Figure 2.**
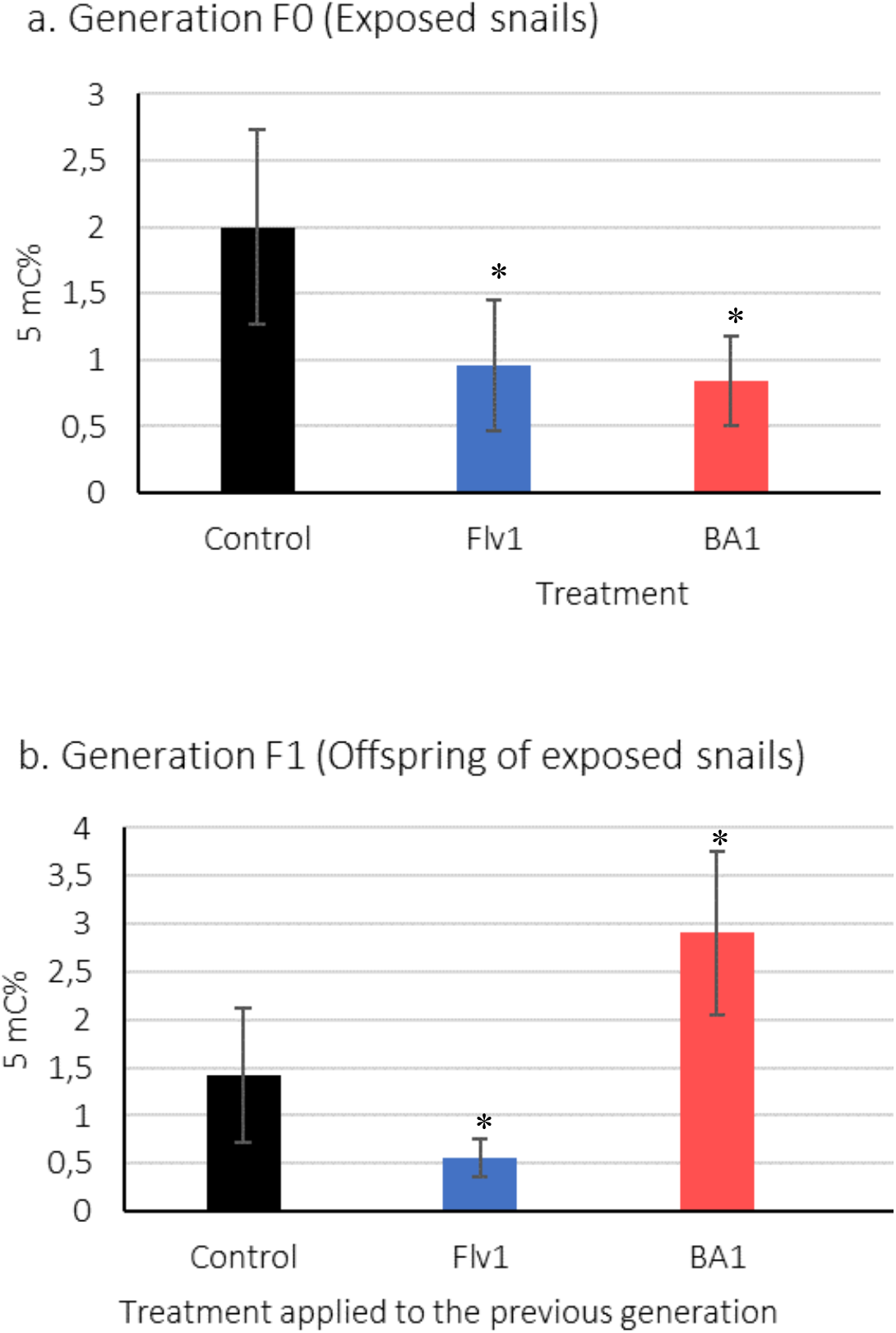
a) Dot blot results showing the 5mC% of the Flv1 and BA1 treated snails compared to control group (F0 generation). B) 5mC% of the F1 generation, offspring of control snails (black bar) and of snails treated with the Flv1 inhibitor (blue bar) and the BA1 inhibitor (red bar). Mann-Whitney Wilcoxon test was applied, significant differences between control and treatment are marked as * for p<0. 05. 5mC ng was normalized to the 5mC % present in the genome of *B. glabrata*.

The results of the ELISA-based detection of 5mC%, showed that the snails paired to produce the epilines presented high global methylation level (highlighted in red in Table 1) in the offspring of BA1-treated snails and low global methylation level (blue in Table 1) in the offspring of Flv1-treated snails and control individuals and some treated snails showed ∼2% of global 5mC% (Table 1). The lines 2, 3, 7, 8, 9, and 12 present the most contrasting profiles of methylation as they showed the highest difference in their 5mC% between paired snails (Figure 3).

**Table 1.** ELISA results showing the 5mC% of the snails of the F1generation (offspring from F0 treated either with Flv1 or BA1) used to generate the epi-lines (F2 generation) and the eight control snails used to generate the four control lines. The snails paired to produce the epi-lines present either low methylation level (highlighted in blue), high methylation level (highlighted in red) or 2% of 5mC%.

| Snail Flv1 (F1, offspring from Flv1 treated snails) | 5mC % | Snail BA1 (F1, offspring from BA1 treated snails) | 5mC% | Gradient in 5mC% in F1 couples (Figure 5) | Line |
| --- | --- | --- | --- | --- | --- |
| Control 1 | 2.11 | Control 1 | 2.16 | 0.05 | Control line 1 |
| Control 2 | 2.0 | Control 2 | 2.02 | 0.02 | Control line 2 |
| Control 3 | 2.20 | Control 3 | 2.10 | 0.10 | Control line 3 |
| Control 4 | 2.13 | Control 4 | 2.06 | 0.07 | Control line 4 |
| Flv1.1-1 | 1.99 | BA1-1 | 3.36 | 1.37 | Epi-line 1 |
| Flv1.1-2 | 0.7 | BA1-2 | 3.05 | 2.35 | Epi-line 2 |
| Flv1.1-3 | 1.8 | BA1-3 | 4.82 | 3 | Epi-line 3 |
| Flv1.1-4 | 2.2 | BA1-4 | 3.35 | 1.15 | Epi-line 4 |
| Flv1.1-5 | 2.31 | BA1-5 | 3.9 | 1.59 | Epi-line 5 |
| Flv1.1-6 | 1.32 | BA1-6 | 1.5 | 0.18 | Epi-line 6 |
| Flv1.1-7 | 1.54 | BA1-7 | 4.82 | 3.28 | Epi-line 7 |
| Flv1.1-8 | 1.16 | BA1-8 | 3.55 | 2.39 | Epi-line 8 |
| Flv1.1-9 | 1.5 | BA1-9 | 3.84 | 2.34 | Epi-line 9 |
| Flv1.1-10 | 1.87 | BA1-10 | 1.97 | 0.1 | Epi-line 10 |
| Flv1.1-11 | 1.83 | BA1-11 | 3.4 | 1.57 | Epi-line 11 |
| Flv1.1-12 | 1.42 | BA1-12 | 4.39 | 2.97 | Epi-line 12 |
| Flv1.1-13 | 1.27 | BA1-13 | 1.36 | 0.09 | Epi-line 13 |
| Flv1.1-14 | 0.31 | BA1-14 | 0.61 | 0.3 | Epi-line 14 |
| Flv1.1-15 | 0.54 | BA1-15 | 1.08 | 0.54 | Epi-line 15 |

After having confirmed that there were differences in DNA methylation between epi-lines we proceeded to phenotyping.

### Epi-lines and epiRILs have higher variation in hatching-rate compared to controls

The most divergent fecundities by cross-fertilization were found in the epi-lines 2, 3, 7, 8, 9 and 12 showing higher hatching rate % compared to controls (Figure 4a).

**Figure 4.**
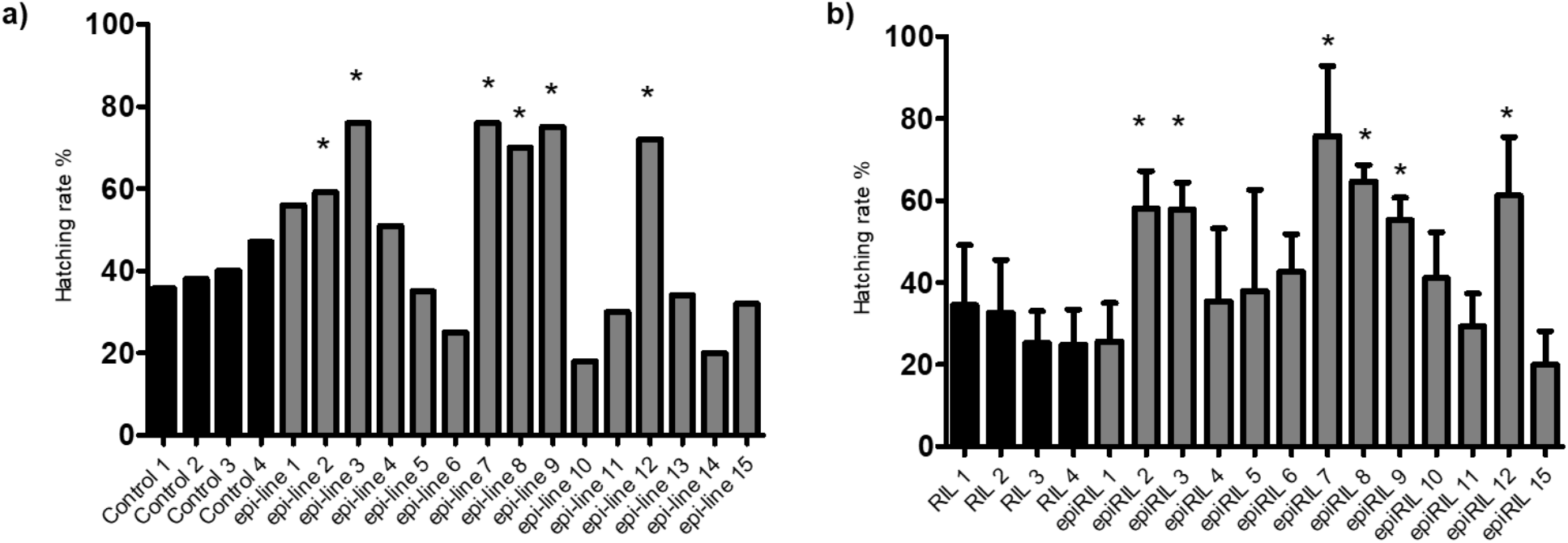
a) Hatching rate of the F2 generation obtained from cross-fertilization of the F1 generation. In black, the hatching rate of control lines. In gray, the hatching rate of the 15 epi-lines. The asterisks (*) indicate significant differences compared to control lines (Fisher’s exact test, p<0.05). b) Hatching rate of the F3 generation obtained from self-fertilization of the F2 generation. The black bars are the RILs, and the gray bars are the epiRILs. The asterisks (*) indicate significant differences compared to controls (One-way ANOVA and Dunnett’s multiple comparisons test, p<0.005). epiRILs 13 and 14 did not reproduce in self-fertilization.

The epiRILs (Fig. 4b) showed more variability in the hatching rate than control RILs. The median of the hatching rate was significantly higher (roughly 2-fold) in 6 out of 13 of the epiRILs (2, 3, 7, 8, 9 and 12) compared to RILs. Interestingly, these epiRILs are offspring of the snails that presented the highest differences in the 5mC% levels.

We calculate the heritability of the fecundity between F2 and F3 generation by calculating the slope of a linear regression. We found that the heritability of the fecundity in the epiRILs varies between 0.5 and0.6 depending on whether or not pseudo replicates in F3 generation are considered (Supplementary file 2, Figures S1-S2 and Tables S4-S5). To simplify our theoretical approach, we decided to use only one value of heritability (*h*^*2*^ = 0.6) for further use in our mathematical model.

To gain further insights into the heritability of fecundity we split individuals of the treatment group (F2 generation) in two groups, those with higher fecundity than controls and those with lower fecundity than controls. We found that in the F2 generation, the values of fecundity of the control group individuals did not overlap with any of the values of the higher or lower fecundity groups (Supplementary file 2, Figure S3). The analysis revealed that in the generation F2, the higher and lower fecundity groups were significantly higher and lower than the control group, respectively (*p* = 0.001 and *p* = 0.04, Supplementary file 2, Table S6). Only one of these two differences is conserved in the F3 generation. Indeed, only the higher fecundity group has a significantly higher fecundity than the control group (*p* = 0.004, Supplementary file 2, Table S7). The lower fecundity group in the F3 generation shows a fecundity that is not different from the control group (*p* =0.46, Table S7). Therefore, the effect of the treatment on fecundity is conserved in the offspring generation, but only in offspring whose parents showed an increased fecundity in response to the treatment.

### epiRILs have higher variation in parasite prevalence compared to controls

The control RILs showed a prevalence ranging from 80 to 90% with a mean of 86%±4 with not significant differences between them (Kruskal-Wallis test, p=0.1). In contrast, the prevalence of the parasite in epiRILs showed significant differences compared to control RILs (Dunnett’s multiple comparisons test, p<0.005). Average prevalence over all epiRILs was decreased to 68±21% with some population’s low extremes (e.g., epiRIL 8 down to ∼20%, and epiRILs 3 and 9 down to ∼30%). (Figure 5).

**Figure 5.**
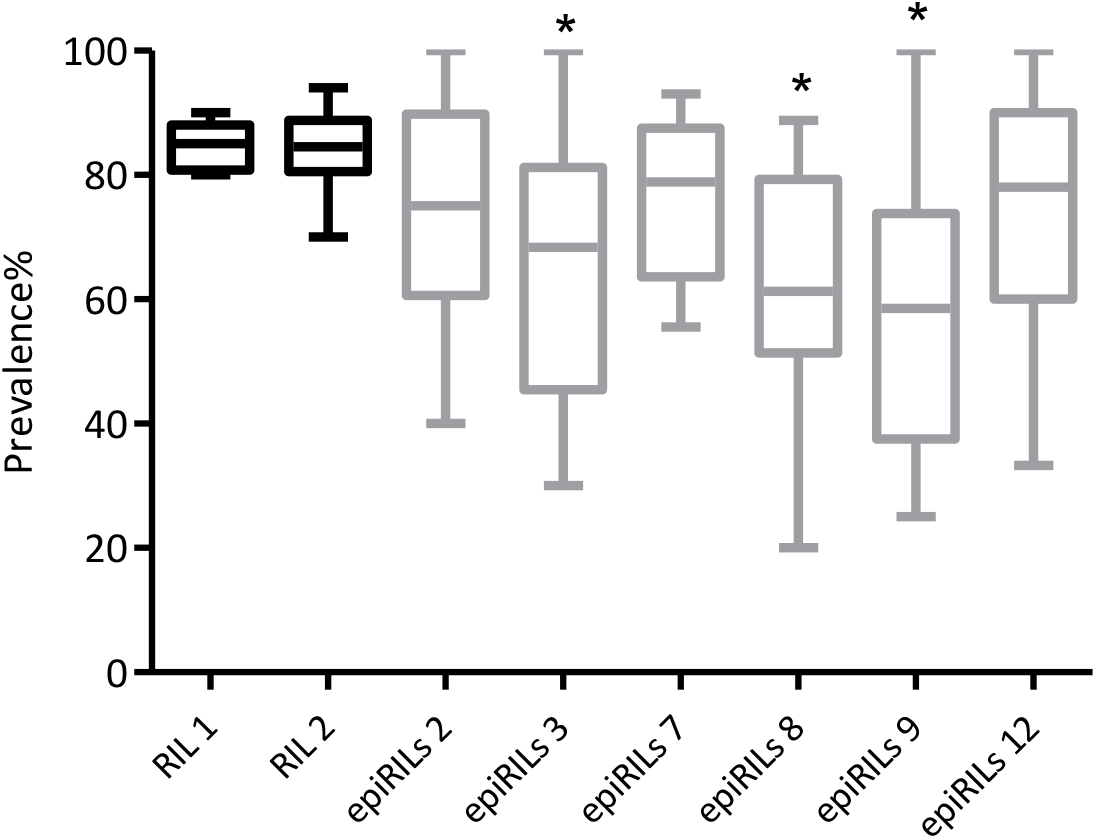
Prevalence of the parasite infestation in control RILs (black boxplots) and epiRILs (gray boxplots) (*n*= 15 per control RIL and epiRIL). The two RILs and the six epiRILs with the highest fecundities in F3 generation were selected to have enough samples for experimental infestations. Data is displayed in Tukey boxplots, representing median and interquartile ranges (IQR). The asterisks (*) indicate significant differences compared to controls (One-way ANOVA and Dunnett’s multiple comparisons test, p<0.05).

The frequency distribution representation of data (Figure 6) shows that the distribution in epiRILs is wider than in control RILs and that new phenotypes occur.

**Figure 6.**
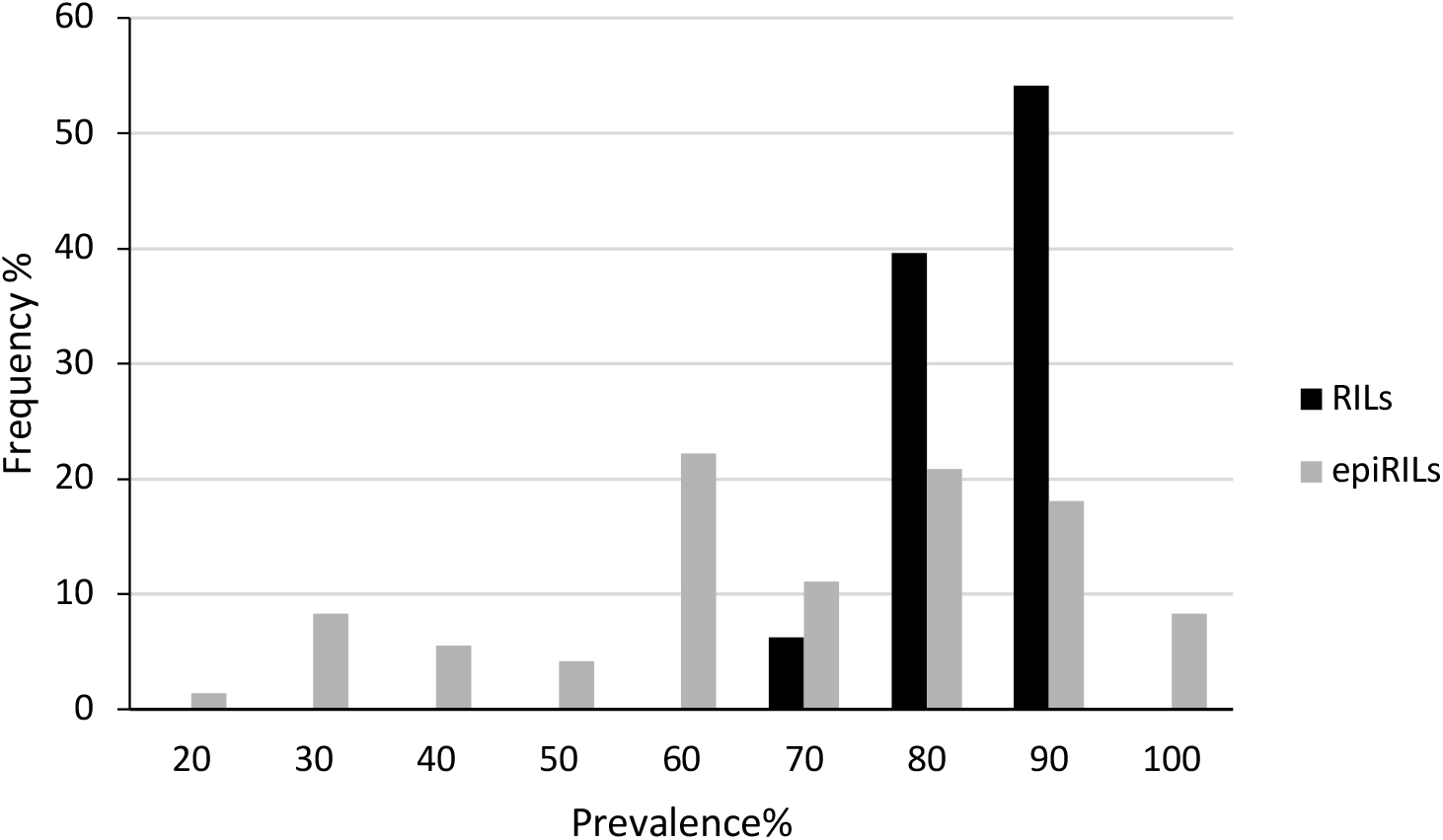
Frequency distribution of prevalence phenotypes in F3. X-axis prevalence in %, y-axis frequency in % in each test population. RILs (n=200), epiRILs (n=600).

### epiRILs have higher variation in parasite intensities of infestation compared to controls

The intensity of infestation is a measure of the number of parasites which penetrate the snails and successfully develop into primary sporocysts. Since the experimental infestations were performed with 5 miracidia (infectious larvae) per snail, the maximum intensity of infestation expected is 5 and the minimum is 0. The intensity of infestation between control RILs was similar (Figure 7), showing a media between 1.93-2.20 and no significant differences were found (Kruskal-Wallis test, p=0.1).

**Figure 7.**
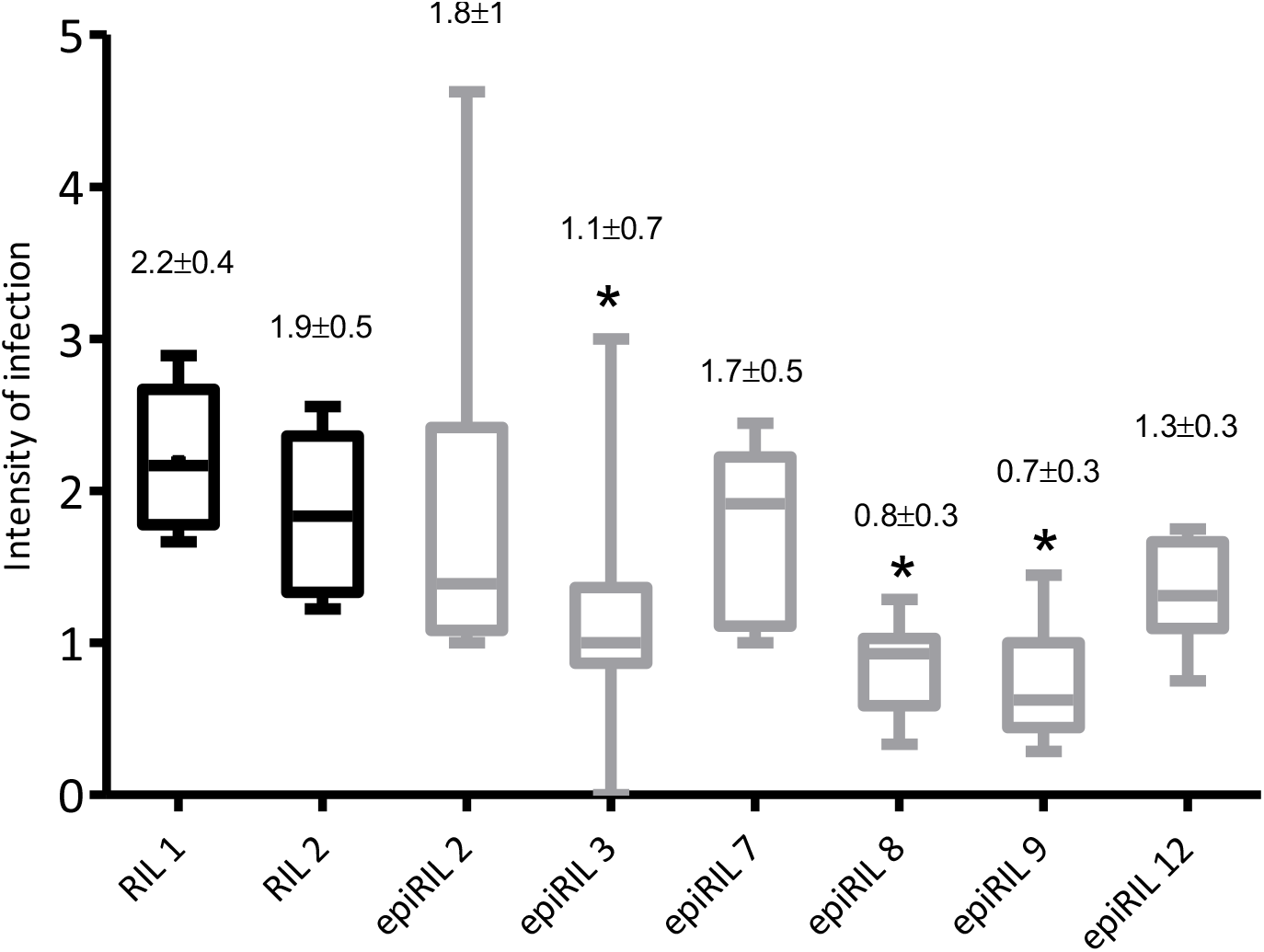
Intensity of infestation in RILs (black boxplots) and epiRILs (gray boxplots). The Kruskal-Wallis and the Dunn’s multiple comparison test were applied. The significant differences against control RILs are marked with * for p< 0.05.

In contrast, we found significant differences against control RILs in the epiRILs 3, 8 and 9 that correspond to the same epiRILs that showed significant differences against control RILs in the prevalence of infestation.

In summary, the lines in which we introduced DNA methylation based epimutations on a homogenous genetic background with a limited variety of alleles showed higher variation in infestation resistance and fecundity. To gain further insights into the nature of DNA methylation changes, we used a new method of reduced representation called epigenotyping by sequencing (epiGBS). Eight individuals from the RIL 1, eight from the epiRIL 8, and eight from the epiRIL 9 were chosen for the analysis of CpG methylation variation and single nucleotide polymorphisms (SNPs) in relation to the contrasted phenotype in prevalence and intensity of infestation. This method was chosen because it represents a cost-efficient alternative to exhaustive methods such as Whole Genome Bisulfite Sequencing similar to Reduced Representation Bisulfite Sequencing that cannot be used in species without CpG islands such *B. glabrata*.

### The epiRILs presented differential methylated cytosines compared to the RILs

After quality control and alignment to the reference genome, the mean percentage of mapping efficiency was 34% in the epiRIL 8, and 35% in the epiRIL 9 and the RIL 1. After methylation calling, we obtained 6 315 CG positions in the epiRIL 8, 27 282 in the epiRIL 9 and 212 031 in the RIL 1.

The epiRILs 8 and 9 that showed significant low prevalence of infestation compared to RIL1 (Figure 7), and displayed significant differences also in their methylation profiles, presenting differential methylated cytosines (DMCs) that were either hypomethylated or hypermethylated.

We found 6 hypomethylated and 33 hypermethylated DMCs between the epiRILs 8 and the RILs. For the epiRIL 9, we found 121 hypomethylated and 201 hypermethylated DMCs compared to the controls (Table S1 and S2, Supplementary file 1). Since DNA methylation differences were introduced by chemical treatment only in the F0 generation, identification of these DMC also demonstrates heritability of epimutations through four generations.

We performed PCA analysis of the CpG methylation in the 1010 cytosine positions for which we had data in all individuals of RIL 1 and epiRIL 9. We found that the groups were more dispersed according to their CpG methylation than according to their SNPs (Figure 8).

**Figure 8.**
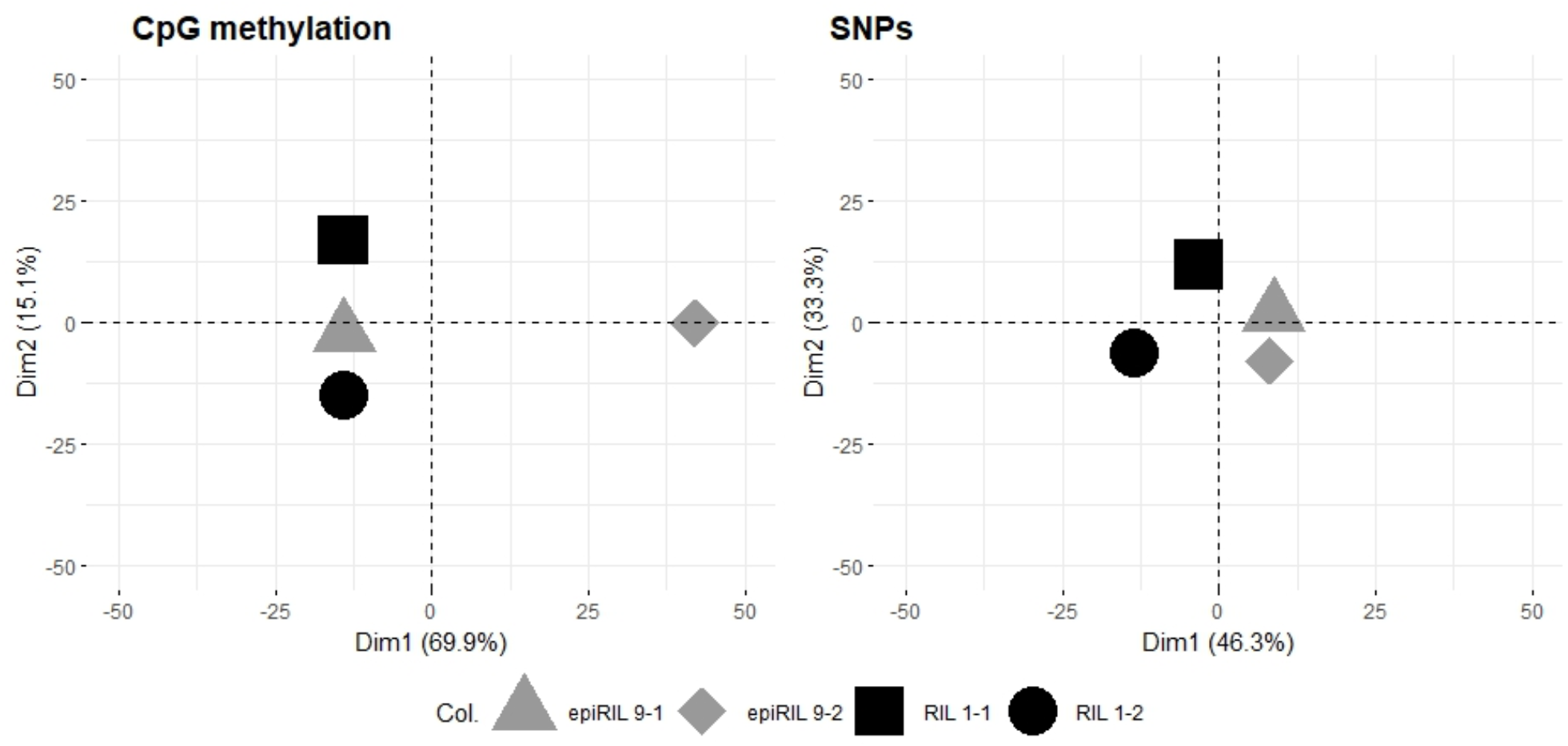
PCA of CpG methylation (left) and SNP variation (right) between the replicates of the two groups RIL 1 and epiRIL 9.

### Mathematical model predicts ephemeral impact of introduction of epimutant offspring unless genetic assimilation occurs

Since we had (i) shown that population-wide prevalence of infestation decreases in offspring of epimutants (Figure 8) and (ii) quantified the heritability of a complex trait to be in a range that is generally considered as “strongly heritable”, we wondered what impact would have the introduction of epimutant offspring into a resident snail community. We fed into the mathematical model also published results that showed a decrease in birth rate and an increase in death rate in infected snails. The epimutant offspring was more resistant and had therefore higher fitness than a resident population, but heritability of their epimutation based traits was less stable (0.6 compared to 1 in the resident population).

Our model predicts that even though the introduced offspring of epimutants would have lower prevalence, their higher fitness would not be sufficient if inheritance would be based exclusively on epigenetic mechanisms with a heritability of 0.6. Even introduction at every generation (roughly every 2 month for *B*.*glabrata*) would gain only marginal decrease in community-wide prevalence. However, if we assume that in a small fraction of the introduced epimutant snails genetic assimilation (*i*.*e*. heritability of 1) of the trait occurred, the scenario would be completely different and would lead to a replacement of the susceptible phenotype by the resistant phenotype after 50-70 generations (Figure 9).

**Figure 9.**
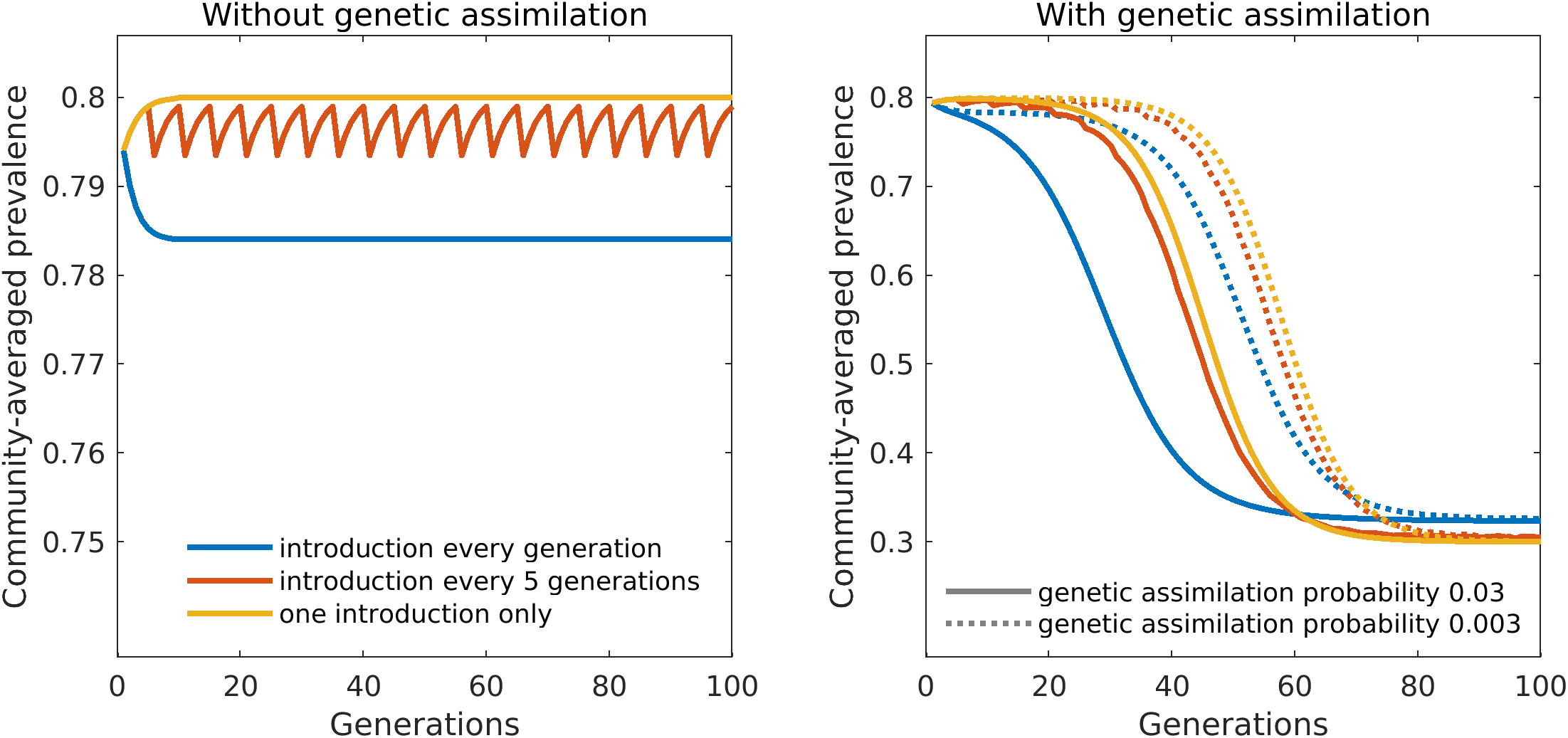
Model predictions for introductions of epimutant snails. We monitor through time the average prevalence of the community consisting of both resident and introduced snails. Left panel: Without genetic assimilation the effect of the introductions is small. Community-averaged prevalence decreases only slightly below *p*_*res*_ = 0.8 (the initial prevalence of the resident snails), even in case of frequently repeated introductions (compare different colors). Right panel: If epimutations can be genetically assimilated, community-averaged prevalence drops from *p*_*res*_ = 0.8 to *p*^*0*^_*mut*_ = 0.3, the prevalence of epimutant snails immediately after introduction. This decrease occurs at later times when genetic assimilation is less probable, after about 40 generations (∼8 years, full lines) or about 60 generations (∼12 years, dotted lines) for the two genetic assimilation probabilities considered here. Note that the panels have a different scale of the y-axis.

## Discussion

In this work, we used Flv1 and BA1 DNMT modulators (DNMTm) on an inbred strain of *B*.*glabrata*, which allowed us to obtain snails with two divergent global epigenomes, one hypo- and one hyper-methylated. By crossing these snails (hyper-paired with hypomethylated) and by subsequent self-fertilization reproduction, we generated epigenetic recombinant inbred lines (epiRILs). We found significant differences in the phenotypes of these epiRILs regarding the fecundity, the prevalence, and the intensity of infestation compared to controls.

We measured two phenotypic traits of importance for parasite transmission: the prevalence and intensity of infestation. The prevalence and intensities of infestation showed high variability in the epi-lines on the contrary to the control lines, which did not show significant differences neither in prevalence nor in intensity of infestation. The sympatric combination used in this study, consisting in *S. mansoni* strain Bresil (*SmBRE*) and the *B. glabrata* strain Bresil (*BgBRE*), is known to display consistent prevalence of infestation without significant variations over multiple generations. This consistency has been observed in our laboratory for at least six generations (Fneich et al., 2016), indicating high heritability of the trait under constant environmental conditions. Indeed, the infestation traits displayed low variability in our control snails. On the contrary, several epiRILs were significantly less infected than control RILs, displaying a significant decrease in the prevalence down to 20% and lower intensities of infestation than control RILs. Not only the prevalence and intensity of infestations were reduced, but the variability of this phenotype increase in the epiRILs. Since we kept genetic variation to a minimum and environmental conditions remained constant, the most parsimonious explanation is that the epiRILs present epigenetic variability resulting into a higher phenotypic variability when exposed to the parasite. Our findings are in line with earlier observations: it is known that infestation with *S. mansoni* induces alteration in the *B. glabrata* DNA methylation, e.g., the expression of DNMT1 and MBD enzymes increased in the snail when exposed to the parasite (Geyer, Niazi et al. 2017) probably altering the transcription of genes involved in the immune response or the response to environmental stress. It has also been found that *S. mansoni* controls the cell spatio-epigenetics of the snail to ensure parasitism. *S. mansoni* induces the relocation of the heat shock protein 70 gene (Bg-HSP70) to the nucleus to be transcribed and hypomethylation of the intergenic region of the gene Bg-HSP70 takes place in temporal concordance with its relocation (Knight et al., 2016). Further investigation will be necessary to elucidate the exact mechanisms by which host snail susceptibility variability can be epigenetically changed and which genomic *loci* are involved.

We also found that the fecundity measured as the hatching rate percentage showed high variability in six epi-lines (F2 generation) that were those that presented the highest differences in the global 5mC level between coupled snails (Figure 3). The same epi-lines showed the highest hatching rates in cross- and self-fertilization (2, 3, 7, 8, 9 and 12) and we also found that only the higher fecundity snails were those that inherited this change to the offspring (Figure S3), suggesting an epi-hybrid-vigor effect similar to the one often observed in genetic hybridization (Geyer et al., 2017).

Our findings indicate that DNA methylation changes in inbred *B*.*glabrata* snails lead to variations in two important phenotypic traits: the resistance to schistosome infestation and fecundity. Our results indicate that more rapid phenotypic changes can be generated through epigenetic changes and that this epimutations can bring phenotypic advantages in traits with an adaptative role. Furthermore, we found that if we select for epimutants with advantageous prevalence phenotypes and that if stability of epialleles transmission occurs, we could use this strategy to reduce prevalence of infestation in wild populations.

Our observation that the introduction of epigenetic variability into genetically homogenous populations leads to new phenotypic variants that can explore the fitness landscape and potentially colonize new fitness maxima corresponds to theoretical models published elsewhere (Pál and Miklós, 1999). It is known that fecundity and survival of infected *B*.*glabrata* snails is lower than those of uninfected snails (Looker and Edges, 1979). We showed here that fecundity of epiRILs increases and that their infestation rate globally decreases but showing also more extreme values (20-100%). We performed a Gedankenexperiment in which we raised epimutant snails in confinement conditions and collected their offspring that we introduced into a resident snail population. We arbitrarily assumed that introduced snails would represent 1% of the resident population. Our mathematical simulation predicts that if genetic assimilation occurred with reasonable probability (0.3 – 3% of the introduced snails) the non-susceptible epimutants would outnumber the control snails after about 10-12 years. The replacement would even occur in case of a single introduction, but it would be faster if the introduction would be done at every generation. It is therefore conceivable that introducing offspring of epigenetically modified parental snails could allow assimilation of the resistant / non-compatible phenotype on focal transmission sites. We did not investigate by which mechanism genetic assimilation could occur.

Nevertheless, to go further and to identify epigenetic and (if any) genetic polymorphic *loci* (or both) that are involved in the phenotypic plasticity of the snail in this trait, it will be necessary to produce high resolution epigenetic and genetic maps of snails with different resistance to infestation. This will allow to evaluate the precise contribution of both information for the observed phenotypic changes. The epigenotyping sequencing method epiGBS has already shown to be efficient to measure DNA methylation changes and single nucleotide polymorphisms in a percentage of the genome (Gawehns et al., 2022; van Gurp et al., 2016) but other more exhaustive methods such as WGBS (Kernaleguen et al., 2018) and ATAC-Seq (Buenrostro et al., 2015) will be more suitable for identifying the involved *loci* with high precision. To gain insights into the evolutionary relevance of the epimutations it will be necessary to conduct longitudinal studies over several generations under real-world conditions.

## Acknowledgements

The authors are grateful to K. Verhoeven (NIOO-KNAW) E. Toulza and J.F. Allienne (IHPE) for support during the optimization of the protocol, and to Ludovic Halby (ETaC) for for the synthesis of the Compound 69.

## Data, scripts, code, and supplementary information availability

Raw data are available online at NCBI SRA PRJNA973008(https://www.ncbi.nlm.nih.gov/bioproject/PRJNA973008).

Scripts and Supplementary information are available online: DOI 10.5281/zenodo.8114402. (https://doi.org/10.5281/zenodo.8114403)

