## Supplementary file 2 for "Epigenetic variation causes heritable variation in complex traits in the mollusk *Biomphalaria glabrata*, vector of the human parasite *Schistosoma mansoni*"

**Heritability estimation**

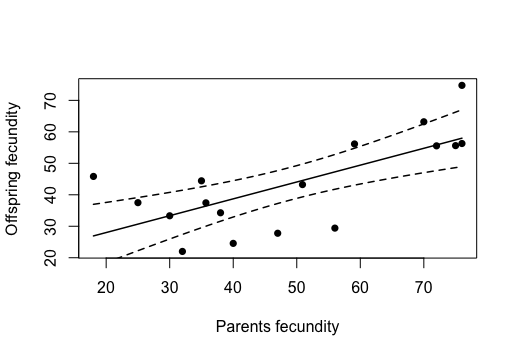

**Figure S1:** Predicted parent-offspring regression with 95% confidence interval. We estimated a heritability of fecundity of 0.53 without pseudoreplicates of the F3 generation (offspring fecundity) (95% CI: 0.23 - 0.83) (Table S4).

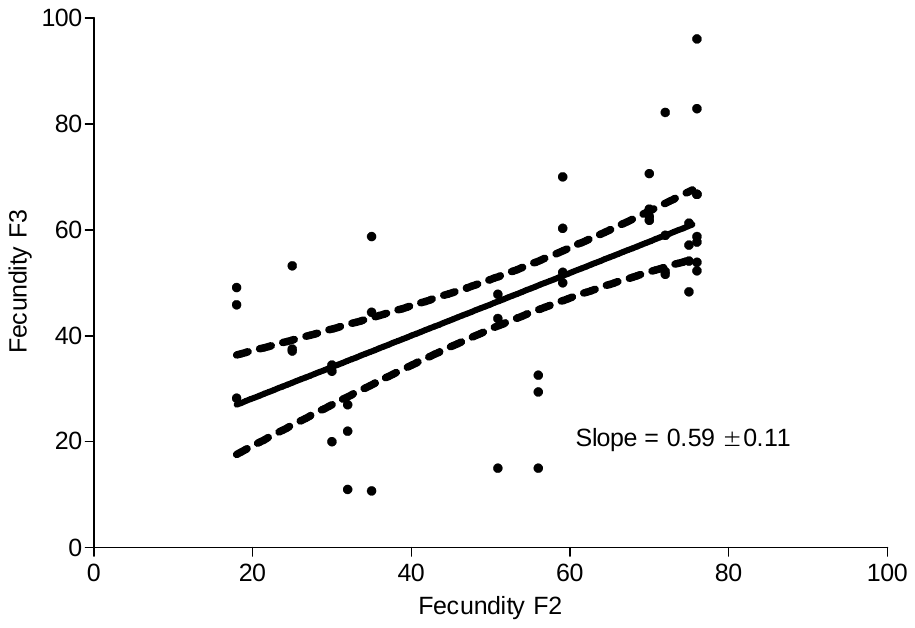

**Figure S2**. Predicted parent-offspring regression with 95% confidence interval. We estimated a heritability of fecundity of 0.59 with the pseudoreplicates of the F3 generation (offspring fecundity) (95% CI: 0.37 - 0.81) (Table S7).

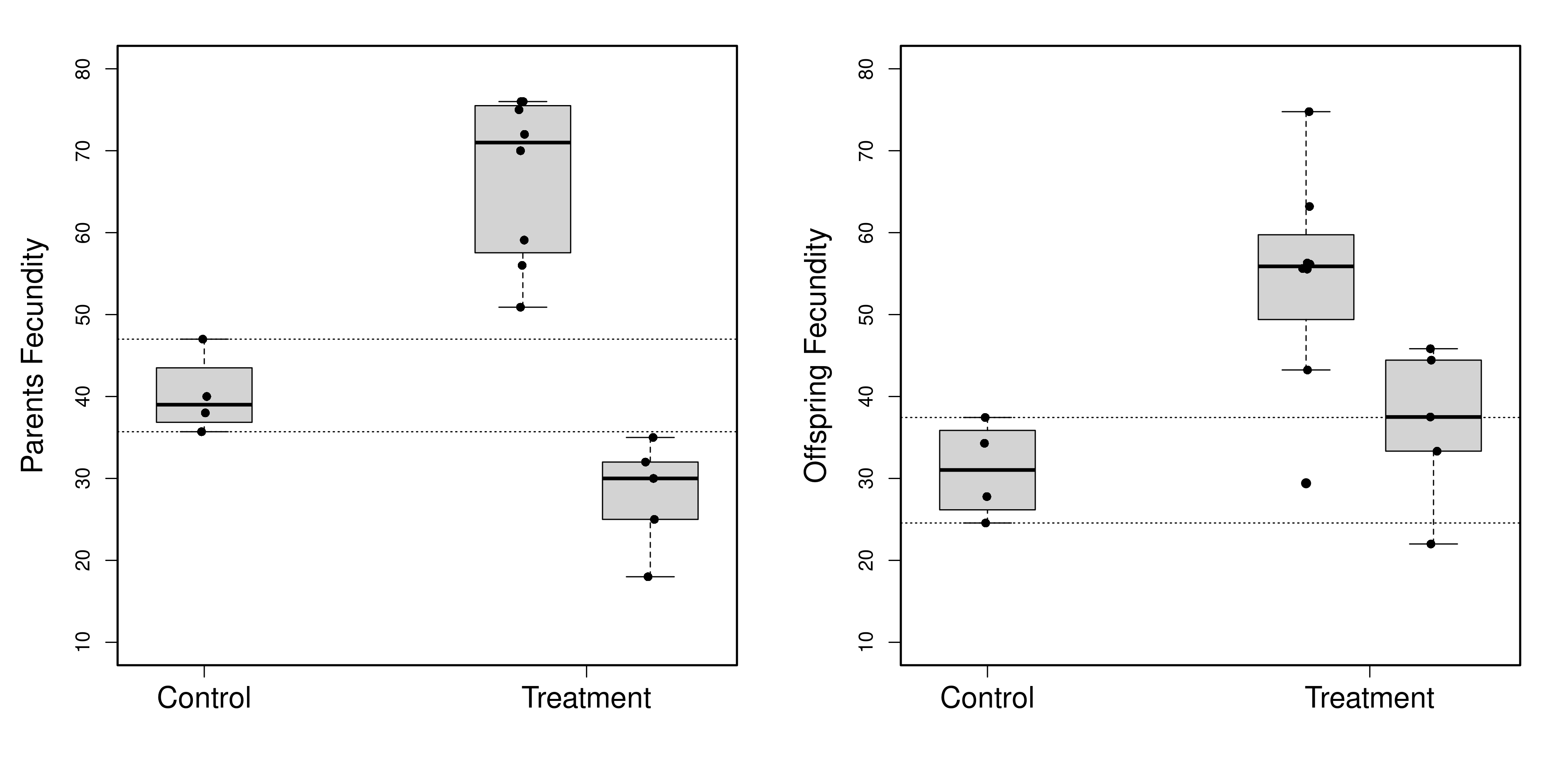

**Figure S3.** Treatment effect on F2 generation (left panel) and F3 generation (right panel)) fecundity. Dots represent the median fecundity value of full sibs. Dotted lines represent the minimum and maximum fecundity values of the F2 generation control group.

**Table S4:** Parent-offspring regression model output

|  | Estimate | Std. Error | t value | p-value |
| --- | --- | --- | --- | --- |
| Intercept | 17.2951 | 7.4061 | 2.335 | 0.03383 |
| Fecundity offspring | 0.5353 | 0.1407 | 3.805 | 0.00173 |

Adjusted R-squared: 0.4572; F-statistic: 14.48, df=15, p-value: 0.001726

**Table S5**. Lineal regression output for F2 generation and F3 generation fecundities with pseudoreplicates in F3 generation (*n* =45).

| Slope | 0.5919 ± 0.1110 |
| --- | --- |
| Y-intercept when X=0.0 | 16.34 ± 6.463 |
| X-intercept when Y=0.0 | -27,60 |
| 1/slope | 1,689 |
| 95% Confidence Intervals |  |
| Slope | 0.3679 to 0.8159 |
| Y-intercept when X=0.0 | 3.296 to 29.38 |
| X-intercept when Y=0.0 | -78.55 to -4.107 |
| Goodness of Fit |  |
| r² | 0,3980 |
| Sy.x | 15,21 |
| Is slope significantly non-zero? |  |
| F | 28,42 |
| DFn, DFd | 1.000, 43.00 |
| P value | < 0.0001 |
| Deviation from zero? | Significant |

**Table S6:** Linear model output for F2 generation fecundity

|  | Estimate | Std. Error | t value | p-value |
| --- | --- | --- | --- | --- |
| Control | 40.17 | 4.12 | 9.74 | 1.3e-07 |
| Group over | 26.70 | 5.05 | 5.29 | 0.00011 |
| Group under | -12.18 | 5.53 | -2.20 | 0.04502 |

Adjusted R^2^: 0.819; F-statistic (2,14) = 37.3, p-value: 2.46e-06

**Table S7:** Linear model output for F3 generation fecundity

|  | Estimate | Std. Error | t value | p-value |
| --- | --- | --- | --- | --- |
| Control | 31.02 | 5.55 | 5.59 | 6.7e-05 |
| Group over | 23.26 | 6.80 | 3.42 | 0.0041 |
| Group under | 5.60 | 7.45 | 0.75 | 0.4643 |

Adjusted R^2^: 0.439; F-statistic (2,14) = 7.26, p-value= 0.00688
